# Imatinib overrides taxane resistance by selective inhibition of novel CLIP1 variant obstructing the microtubule pore

**DOI:** 10.1101/838334

**Authors:** Katsuhiro Kita, Prashant V. Thakkar, Giuseppe Galletti, Neel Madhukar, Elena Vila Navarro, Isabel Barasoain, Holly V. Goodson, Dan Sackett, José Fernando Díaz, Olivier Elemento, Manish A. Shah, Paraskevi Giannakakou

**Author notes:** Correspondence and requests for materials should be addressed to: Paraskevi Giannakakou; Manish A. Shah; Olivier Elemento. These authors have contributed equally to this manuscript (Katsuhiro Kita, Prashant V. Thakkar).

## Abstract

Despite its widespread use, the majority of patients with gastric cancer (GC) will not respond to taxane chemotherapy due to resistance mechanisms. Here, we report the discovery of a novel truncated variant of the microtubule plus-end binding protein (+TIP) CLIP-170, hereafter CLIP-170S, whose expression is enriched in taxane resistant cell lines and patients with GC. To establish causation, we knocked-down (KD) CLIP-170S which completely reversed taxane resistance. Mass-spec proteomics and 5’-RACE further showed that CLIP-170S lacked the first 150 amino acids, including the Cap-Gly motif required for microtubule (MT) plus-end localization. Mechanistically, we show that CLIP-170S was mislocalized from the MT plus-end to the MT lattice obstructing the MT pore surface site required for taxane entry into the MT lumen. Computational analysis of RNA-seq data from taxane-sensitive and resistant GC cell lines, predicted imatinib as the top candidate drug to overcome drug resistance. Imatinib treatment completely reversed taxane resistance, as predicted, and did so unexpectedly by selective depletion of CLIP-170S. Importantly, CLIP170S was found to be highly prevalent in tumor biopsies from patients with GC. Taken together, these data identify CLIP-170S as a novel, clinically prevalent +TIP variant that obstructs the MT pore and confers taxane resistance. The discovery of this previously unrecognized variant together with the computational discovery of Imatinib as a selective CLIP-170S inhibitor, implicate the MT pore in clinical taxane resistance and provide new therapeutic opportunities for treatment of GC and beyond.

## Background

Gastric cancer (GC) is a highly morbid and prevalent global disease and is the second leading cause of cancer related deaths worldwide in 2018^1^. Such a dismal prognosis is a result of advanced stage at diagnosis coupled with demonstrated resistance to chemotherapy^2^ and few available targeted options^3^. Docetaxel has been FDA approved as a first line therapy for GC when combined with Cisplatin and 5–Fluorouracil^4^, and paclitaxel is a standard second line treatment option^5^. Notably, a large proportion of patients demonstrate resistance to taxane treatment, with response rates to single agent therapy around 14-25%, and duration of response measured in months^6,7^. This is further recapitulated in our post-hoc analysis of the trial that led to the approval of docetaxel as first line treatment in GC^8^, in which we identified a significant subset of GC patients who did not derive any clinical benefit from docetaxel chemotherapy. Thus, there is a dire need to decipher clinically actionable mechanisms of taxane resistance.

Despite the multitude of taxane resistance mechanisms described in preclinical models by us and others, including tubulin mutations, altered MT dynamics or the presence of drug efflux mechanisms, none has impacted clinical care^9-19^.

Here we describe a novel truncated variant of the MT plus-end binding protein CLIP170, hereafter CLIP170S, which we found to be significantly enriched in taxane-resistant GC cell lines and patient samples. CLIP-170 binds to the plus-end of growing MTs and is involved in the regulation of MT dynamics, positioning of MT arrays and signaling ^20^. We demonstrate a direct role for CLIP-170S in conferring taxane resistance in gastric cancer by obstruction of the MT pore, thereby preventing taxane internalization into the MT lumen. Using a systems biology approach, we discovered the tyrosine kinase inhibitor, imatinib, as a compound predicted to reverse taxane resistance. Mechanistically we show that imatinib sensitizes resistant GC cell lines to taxane therapy by selective depletion of CLIP-170S. Taken together, this study describes a novel +TIP protein variant with distinct biological properties, a previously unrecognized mechanism of taxane resistance that involves obstruction of the MT pore, and identifies an actionable therapeutic strategy to treat patients with taxane refractory GC.

## Results

### Taxane-resistant GC cells exhibit reduced Flutax-2 residence time on microtubules

We have previously reported that taxane resistance in gastric cancer (GC) is associated with reduced drug target engagement in both cell lines and patient biopsies^8^. Using a panel of 12 GC cell lines, we identified a subset (6/12) with intrinsic resistance to DTX which was due to lack of drug-target engagement, despite unimpaired intracellular drug accumulation and absence of tubulin mutations or other target alterations. To understand the molecular basis of the impaired interaction between taxanes and MTs we incubated native cytoskeletons from sensitive and resistant GC cells with FITC-conjugated paclitaxel (Flutax-2) and determined its binding equilibrium with MTs by live cell imaging, as previously described^21^ (Figure 1A). As can be seen, initially Flutax-2 bound to both sensitive and resistant cell cytoskeletons (Figure 1B, Supplementary figure 1A) and was allowed to equilibrate following wash out. Flutax-2 remained bound to native cytoskeletons from the TMK1 taxane-sensitive cell line for the duration of the experiment (180 min) (Figure 1B). In the case of the resistant Hs746T and SCH cells, the drug had to dissociate significantly in order to reach the binding equilibrium, thus, suggesting a higher dissociation constant for the resistant cell cytoskeletons (Figure 1B and Supplementary Figure 1A).

**Figure 1:**
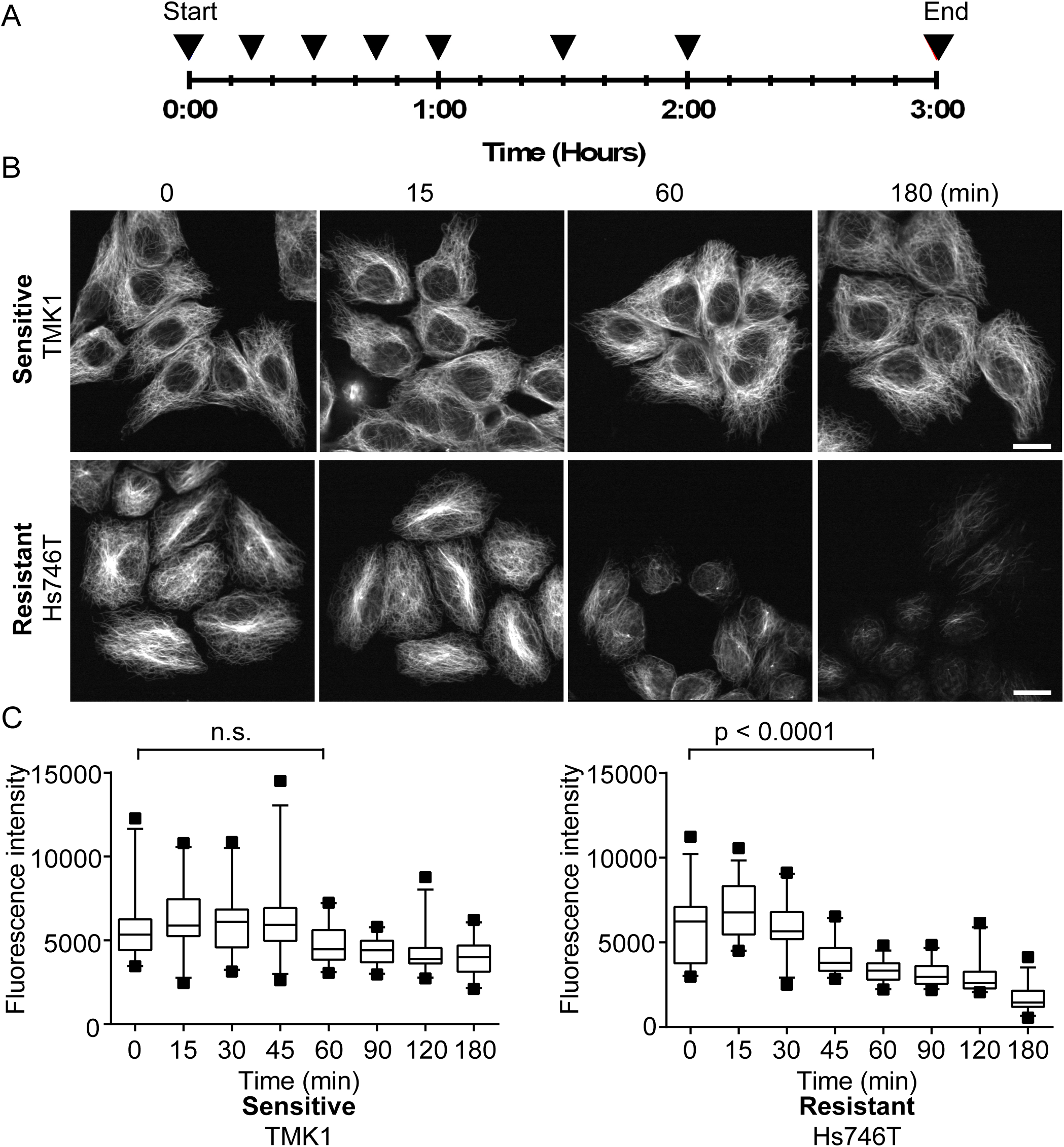
Taxane-resistant GC cells exhibit reduced Flutax-2 residence time on MTs. Native cytoskeletons were prepared from drug sensitive and resistant cell lines and treated with FITC-conjugated paclitaxel (Flutax-2) for 10 min. Following drug washout (0 min), cells were imaged with a spinning disk confocal microscope at 15 min intervals (black arrowheads) for a total of 3 hr A) Schematic diagram of image acquisition. B) Representative images of Flutax-2-labeled native cytoskeletons from DTX-sensitive cell line TMK1 and DTX-resistant cell line Hs746T. bar = 20 µm. C) Box-plot representation of Flutax-2 fluorescence intensity in TMK1 and Hs746T cell line. 5-95% confidence intervals graphs are shown (n=40-70 individual cells/time point/cell line); statistical values for each cell line between 0 and 60 min are shown; Mann-Whitney test; n.s.; not significant.

The equilibrium of Flutax-2 binding is a function of its association rate constant (*k*_*on*_), a bimolecular reaction that depends on the diffusion constant of the drug, and dissociation rate constant (*k*_*off*_), a monomolecular reaction. Thus, an altered *k*_*on*_ in the absence of an altered *k*_*off*_ would imply altered diffusion of the drug to the luminal site of MTs, while an altered *k*_*off*_ alone would imply that the binding site itself is altered.

Quantitation of Flutax-2 association and dissociation rate constants revealed a 3-fold lower *k*_*on*_ value in resistant Hs746T (2 ± 1 × 10^4^ M^−1^ s^−1^) *versus* sensitive TMK1 (6 ± 2 × 10^4^ M^−1^ s^−1^) cells (Supplementary figure 2A). In contrast, no major differences were observed in their respective *k*_*off*_ dvalues (Hs746T: 4.3 ± 0.5×10^−2^ s^−1^ *vs* TMK1: 3.5 ± 0.4 × 10^−2^ s^−1^) (Supplementary figure 2B).

**Figure 2:**
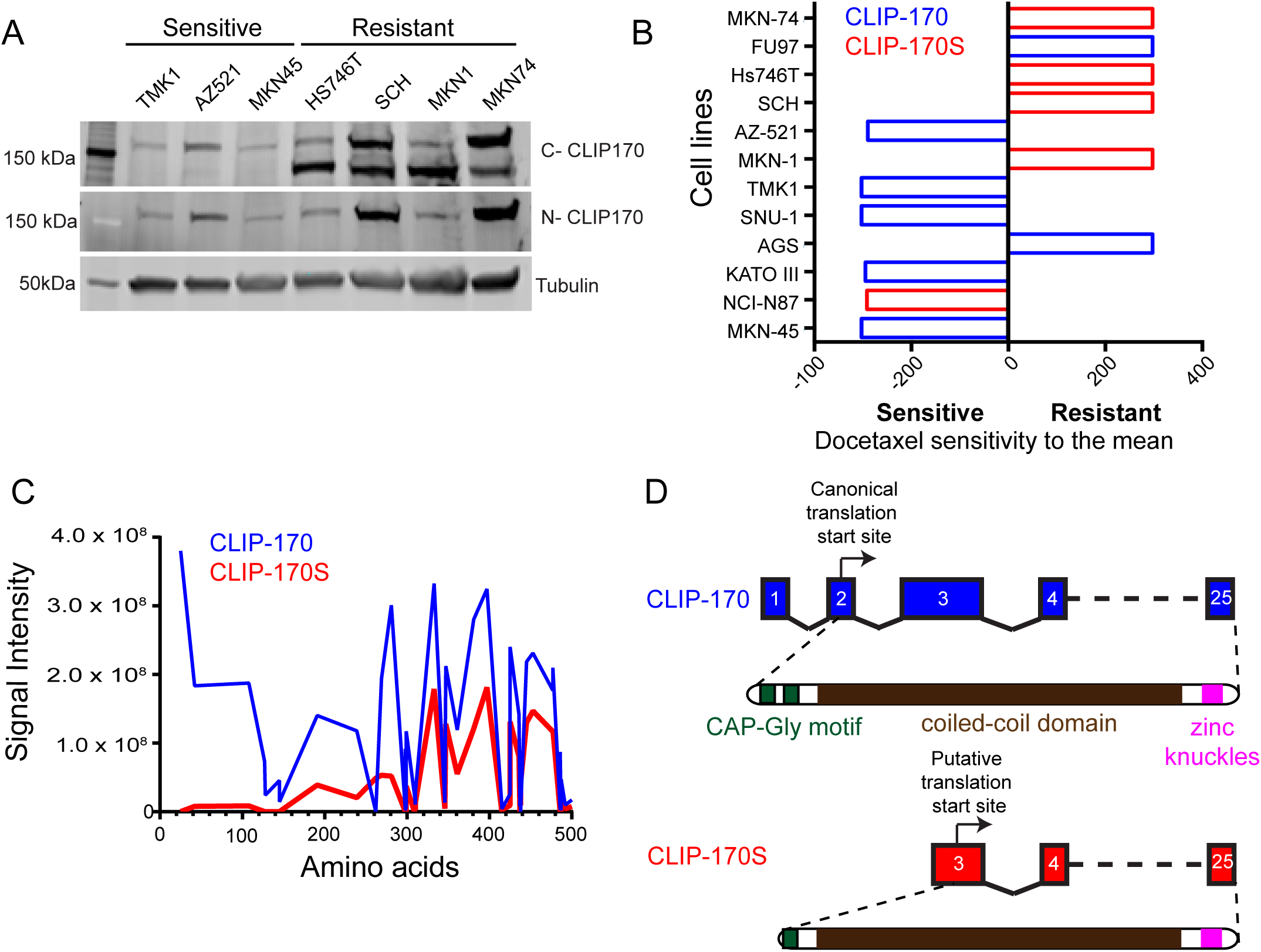
Expression of novel truncated CLIP-170 variant, CLIP-170S, enriched in taxane-resistant GC cell lines. A) CLIP170 expression was assessed by immunoblot in a panel of 7 GC cell lines with known sensitivity/resistance to DTX. Immunoblot was performed using antibodies against the C-terminus or N-terminus of CLIP-170 or tubulin, as indicated. Immunoblot with anti C-CLIP170 antibody revealed the presence of a faster migrating band in DTX-resistant cell lines. B) Mean Graph representation of relative sensitivity/resistance to DTX in an expanded panel of 12 GC cell lines. The mean IC_50_ value of all cell lines is calculated and set at 0. Individual IC_50_ values are shown as bars projecting in opposite directions based on whether cell sensitivity is higher (relative resistance, red bars) or lower (relative sensitivity, blue bars) than the mean. CLIP170 immunoblot was performed in the 12 cell lines as in A and CLIP-170 expression status was superimposed to the Mean Graph. Blue: canonical only CLIP-170; Red: co-expression of CLIP-170S. CLIP-170S expression is significantly enriched in the DTX-resistant cell lines, p<0.05; one-tailed Student’s t-test, data are representative of three individual biological repeats. C) Immunoprecipitation using the anti C-CLIP170 antibody followed by mass spectrometric analysis revealed that CLIP-170S lacks the first N-terminal 155 amino acids confirming the size and identity of the faster migrating protein in A, as a truncated CLIP-170 variant with a predicted molecular weight of 152kDa. D) Schematic representation of exon and protein structures of canonical CLIP-170 and CLIP-170S as identified by 5’-RACE and mass spectrometry. CLIP-170S by missing the first 1∼155 amino acids lacks the first Cap-Gly domain that mediates proper plus-end localization. Arrows indicate position of canonical translation start site for CLIP-170 and putative translation start site for CLIP-170S.

Taken together, these data demonstrate that the lower equilibrium binding constant of taxanes for resistant cell cytoskeletons arises from the slower association rate constant (*k*_*on*_) indicating an impaired diffusion of the drug to the luminal binding site in MTs ^22^. Our proposed mechanism suggests a paradigm-shift in our understanding of taxane resistance, wherein impaired drug access to the binding site, as opposed to site structural alterations, affects target engagement.

### Identification of a novel N-terminal truncated variant of plus-end binding protein CLIP-170

In characterizing sensitive and resistant GC cell lines, we observed altered microtubule dynamics at baseline, quantified by live cell imaging of MT plus ends labeled by EB1-GFP. We found that resistant cells displayed significantly lower rates of MT growth speed and rescue/repolymerization as compared to the sensitive GC cell lines^8^, suggesting that MT dynamics and the proteins regulating them may underlie taxane resistance in these cells. We focused particularly on one such MT +TIP protein, the CAP-Gly domain containing linker protein, CLIP-170, expression of which has been shown to have an inverse correlation with taxane resistance in breast cancer cell lines^23^. CLIP-170 was the first identified +TIP protein that binds to MT plus-ends via its association with End-binding protein-1 (EB1) and links MTs to endoplasmic vesicles in addition to regulating MT dynamics ^24-28^ (reviewed in ^20,29^).

Interrogation of GC sensitive and resistant lines for CLIP-170 protein expression revealed the presence of a novel faster migrating isoform of CLIP-170, hereafter CLIP-170S, in addition to the canonical full-length protein (Figure 2A). Notably, this result was obtained when we used an anti-C-terminal CLIP-170 antibody, while anti-N-terminal antibody detected only the canonical protein, suggesting an N-terminal truncation in CLIP-170S. Interestingly, CLIP-170S was preferentially expressed in taxane-resistant GC cell lines (Figure 2A). Expanding this analysis to the entire panel of 12 GC cell lines revealed that CLIP-170S was significantly enriched in taxane-resistant cell lines with 4/6 resistant *versus* only 1/6 sensitive cell lines expressing CLIP-170S (p< 0.05) (Figure 2B). We next performed immunoprecipitation followed by mass spectrometry and showed that CLIP-170S lacks the first 150 amino acids from its N-terminus compared to the canonical full-length protein (Figure 2C), confirming the size and identity of the faster migrating protein in 2A. Consistent with these results, 5’-rapid amplification of cDNA ends (5’-RACE) identified two CLIP-170 transcripts in the Hs746T and SCH resistant cells expressing both variants. The first transcript corresponded to the canonical CLIP-170 while the second, less abundant transcript, had a start site in exon 3, in the resistant but not in the sensitive GC lines (Supplementary figure 3). These transcripts that start in exon 3 would utilize the putative start codon at position 620 (amino acid position 155) and therefore, generate a protein of about 140kDa in agreement with the results in 2A. Based on these results we generated a predicted exon structure for CLIP-170S with the putative translation start site and the known protein domains of CLIP-170 displayed (Figure 2D). Taken together, we have identified CLIP-170S as a novel, shorter protein variant of CLIP-170, expression of which is preferentially enriched in taxane resistant GC cell lines.

**Figure 3:**
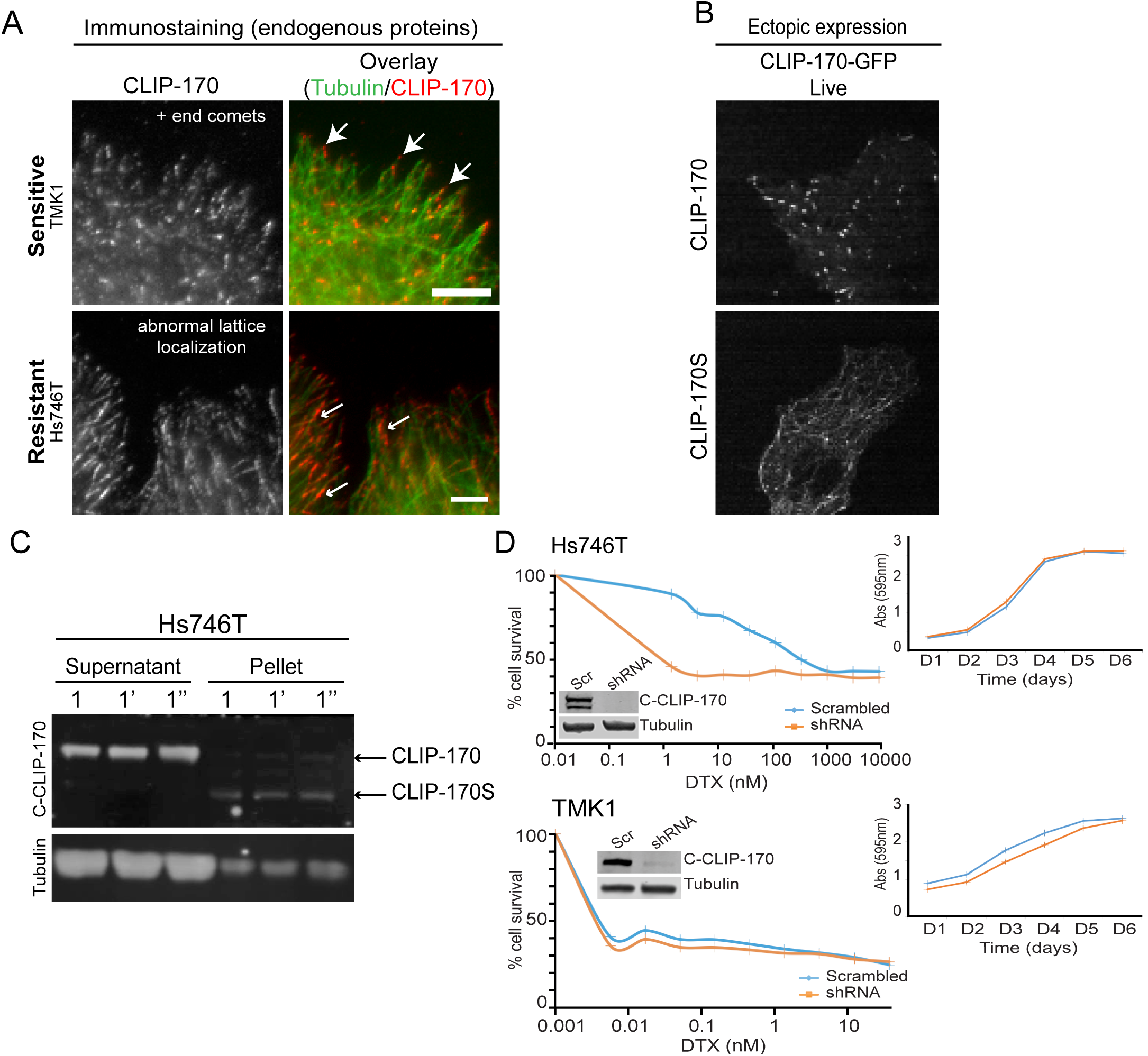
CLIP-170S is mislocalized from the MT plus-end to the lattice, and its knock-down sensitizes cells to taxane. A) CLIP-170 cellular distribution was assessed by confocal microscopy following immunofluorescence staining of endogenous proteins using antibodies against the C-terminus of CLIP-170 or tubulin in sensitive and resistant cell lines. MT plus-end CLIP-170 localization (large arrows) is observed in the TMK1 cells expressing canonical only CLIP-170; CLIP-170 lattice localization (small arrows) is observed in the Hs746T cells expressing both canonical and CLIP-170S. B) Ectopic expression of GFP-tagged CLIP-170 and CLIP-170S in COS-7 cells. Single frames were extracted following live cell imaging from cells expressing similar low levels of each protein. Notice the comet-like pattern of canonical CLIP-170 in contrast to the MT lattice distribution pattern exhibited by CLIP-170S. C) MT polymers were separated from soluble tubulin dimers from Hs746T cells following centrifugation into pellet and supernatant fractions, respectively, and immunoblotted using antibodies against the C-terminus of CLIP-170 (top) or Tubulin (bottom). Lysates from three independent biological repeats (1, 1’, 1”) show reproducible and preferential association of CLIP-170S with MT polymers in the pellet fraction. D) CLIP-170S knockdown sensitizes Hs746T cells to taxanes. Hs746T cells expressing endogenous CLIP-170 and CLIP-170S and TMK1 cells expressing only CLIP-170, were stably knocked-down for CLIP-170 and were plated for cytotoxicity to DTX or for growth rate assessment. Knockdown (KD) of CLIP-170 and CLIP-170S sensitized the resistant HS746T cells by 300-fold (top; IC_50_: Scr = 328 nM, shRNA = 1.3 nM), while CLIP-170-KD had no effect on senstitive TMK1 cells as compared to their respective scrambled controls (bottom; IC_50_: Scr = 0.004 nM, shRNA = 0.004 nM). Representative data from one of several independent biological repeats is shown. Inset shows confirmation of knockdown by immunoblot. Top right insets: growth rates of HS746T and TMK1 cells remain unchanged following CLIP-170-KD as compared to their respective scrambled controls.

### CLIP-170S is mislocalized from the microtubule plus-ends to the lattice

The N-terminus of CLIP-170 contains two cytoskeleton-associated protein glycine-rich (CAP-Gly) domains that mediate MT binding and plus-end tracking via binding to End-binding protein-1 (EB1)^24,25^. Of the two domains the first Cap-Gly binds stronger to EB1, an interaction that is required for plus end tracking, while the second Cap-Gly motif demonstrates higher affinity for MTs ^30-32^.

The predicted exon structure for CLIP-170S indicates loss of the first Cap-Gly motif, suggesting potential defects with plus-end tracking. Immunofluorescence staining of endogenous CLIP170 in taxane sensitive and resistant cells revealed preferential CLIP-170 localization to the MT lattice in cells expressing CLIP-170S, in contrast to the comet like distribution at MT plus-end characteristic of canonical CLIP-170 ^25^ (Figure 3A). Live cell imaging and immunofluorescent staining of exogenously expressed GFP-tagged fusion proteins, also demonstrated that while GFP-CLIP-170S was distinctly localized along the MT lattice, canonical GFP-CLIP170 exhibited the expected comet like localization at MT plus-end, a pattern indistinguishable from staining of endogenous proteins (Figure 3B and Supplementary Figure 4 and Supplementary movies 1 and 2). Additionally, separation of endogenous MT polymers from soluble tubulin dimers by centrifugation in Hs746T cells followed by immunoblot, showed that CLIP-170S preferentially associated with the MT polymers present in the pellet fraction in contrast to CLIP-170 which was predominantly found in the soluble fraction (Figure 3C). Taken together, these data provide evidence for distinct cellular localization of CLIP-170S, preferentially associated with the MT polymers along the lattice and defective plus-end MT tracking.

**Figure 4:**
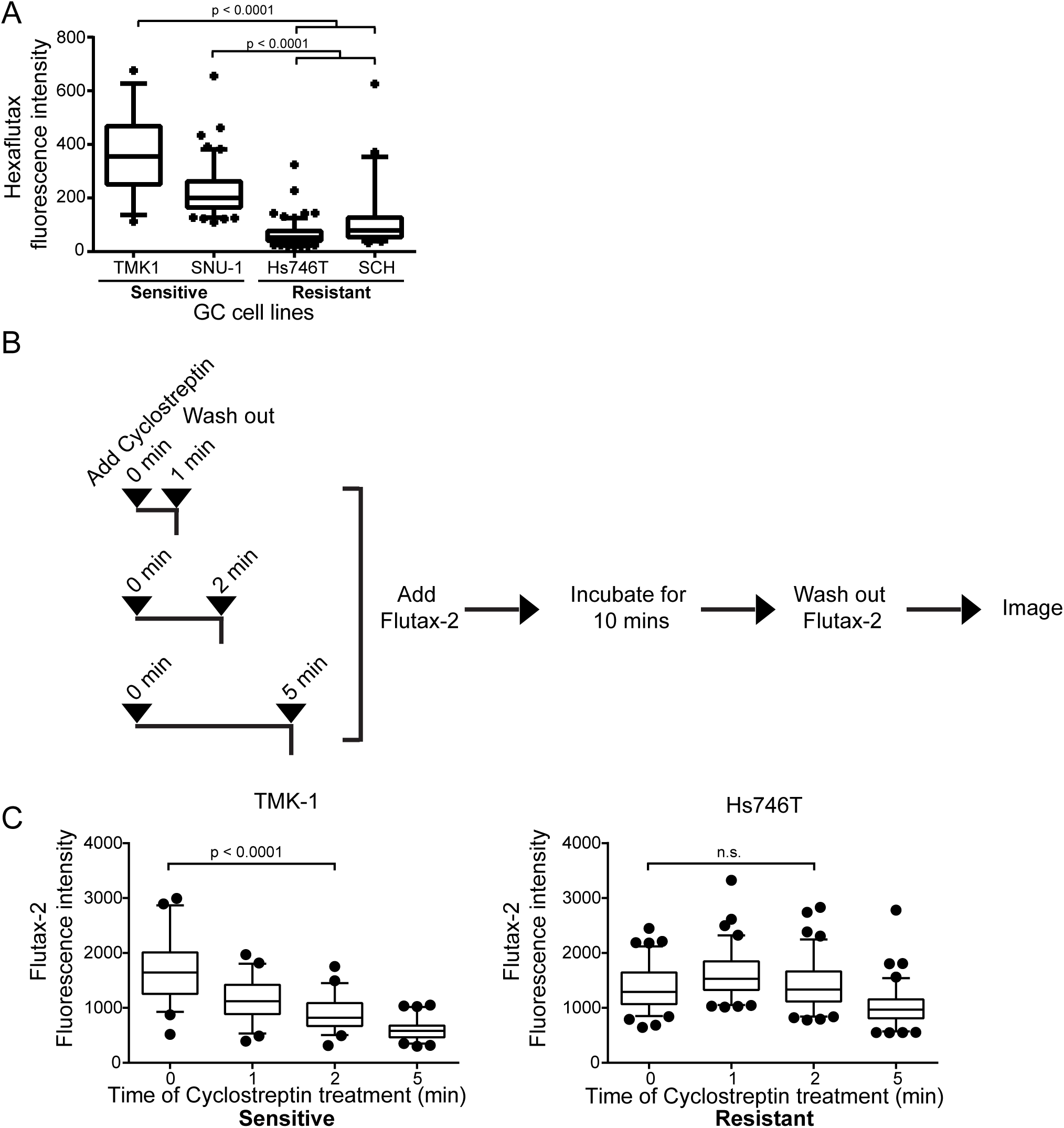
Probing the MT pore using fluorescent taxoids reveals that taxane binding to the outer wall of MTs is partially obscured in resistant GC cells. A) Fluorescent Intensity of Hexaflutax binding to native cytoskeletons in sensitive and resistant GC cell lines. Hexaflutax binds at the outer only surface of the MT pore. Sensitive cell lines TMK1 and SNU-1 display higher hexaflutax binding as compared to the resistant GC cells, Mann-Whitney test. Results are representative of four individual biological repeats. 5-95% confidence interval shown (n = 35-145 cells/timepoint/cell line). B) Schematic illustration of Flutax-2 binding following cyclostreptin pre-treatment of native cytoskeletons. Cells were pre-treated with cyclostreptin for 0, 1, 2 or 5 min, prior to Flutax-2 treatment for 10 min. Native cytoskeletons were then imaged using a spinning disk microscope. C) Box-plot representation of Flutax-2 binding to native cytoskeletons pre-incubated with cyclostreptin. Significant loss in Flutax-2 binding was observed following a 2 min cyclostreptin pre-incubation in TMK1 (left) but not Hs746T cells (right), Mann-Whitney test, 5-95% confidence interval shown (n=100-200 cells/time point/cell line).

### CLIP-170S depletion reverses taxane resistance in GC cells

To determine causation, we stably knocked-down CLIP-170 and CLIP-170S from sensitive and resistant GC cell lines and performed cytotoxicity experiments. Stable knock-down (KD) of both CLIP-170 and CLIP-170S completely reversed taxane resistance (∼300-fold) in Hs 746T cells, whereas knockdown of canonical CLIP-170 only in sensitive TMK1 cells had no effect on taxane sensitivity (Figure 3D). We found no difference in the cellular growth rates in cells expressing scramble or the CLIP-specific shRNA (Figure 3D inset), ruling out faster growth rates as a contributing factor to enhanced taxane sensitivity.

To determine the effect of CLIP-170S depletion on MT residence time of Flutax-2, we treated native cytoskeletons from Hs746T cells stably expressing scrambled or CLIP-170 shRNA with Flutax-2 and quantified drug binding over time. Similar to the results shown in Figure 1C, we observed significant reduction in Flutax-2 binding to native cytoskeletons in control Hs746T-Scrambled shRNA cells (0 min vs 45 mins, p < 0.0001). In contrast, in Hs746T-CLIP-170-KD cells, Flutax-2 binding equilibrium remained unchanged for the duration of the experiment suggesting that CLIP-170 depletion restores Flutax-2 binding to MTs (Supplementary Figure 5). CLIP-170-KD in TMK1 cells expressing only the canonical protein did not have any effect of Flutax-2 binding equilibrium (data not shown). Taken together, these results demonstrate causation between CLIP-170S expression and altered taxane binding efficiency.

**Figure 5:**
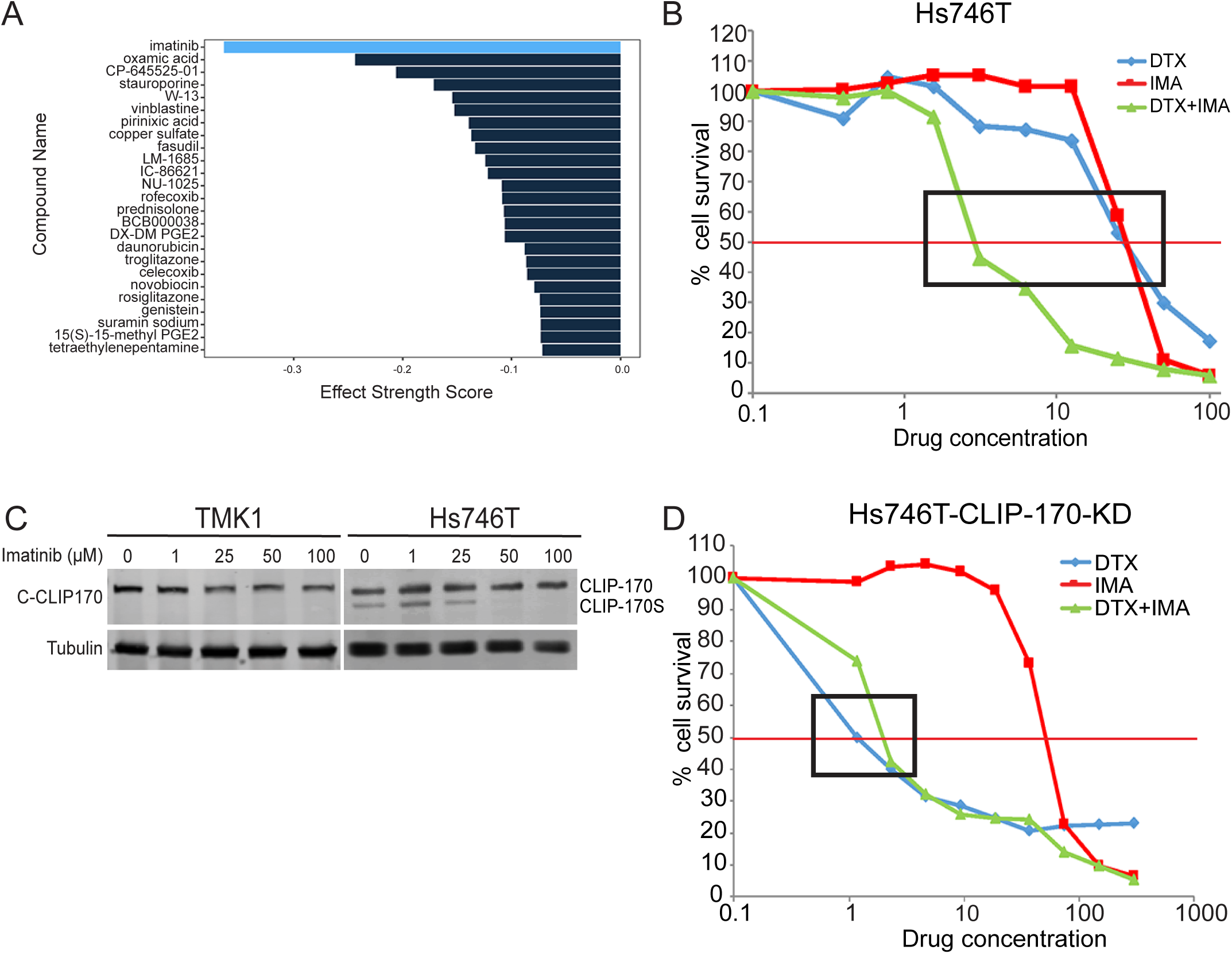
Imatinib reverses taxane resistance in GC by specific depletion of CLIP-170S. A) Gene-expression based computational analysis of RNA-Seq data from untreated and DTX-treated sensitive and resistant GC cell lines were used to derive a DTX-resistance signature. Using the Connectivity Map (CMAP) database, we identified Imatinib as the top candidate to reverse taxane resistance. Bars with more negative values indicate higher likelihood for a compound to reverse DTX resistance. B) Imatinib sensitizes HS746T cells to docetaxel. Cytotoxicity assay using DTX-resistant Hs746T cells treated with docetaxel (DTX) or imatinib (IMA) either alone or in combination (DTX + IMA). C) Imatinib selectively inhibits CLIP-170S expression without affecting CLIP-170. Hs746T and TMK1 cells were treated with 1, 25, 50 and 100 µM Imatinib for 3 hr and processed for immunoblotting using antibodies against the C-terminus of CLIP-170 or tubulin. D) Imatinib has no effect on Hs746T-CLIP-170-KD sensitivity to docetaxel. Cytotoxicity assay as in B.

### CLIP-170S confers taxane resistance via obstruction of the MT pore

To elucidate the mechanism underlying CLIP-170S mediated taxane resistance we focused our investigation on taxane binding to the outer and inner MT surface as CLIP-170S exhibited extensive and aberrant localization to the MT lattice. Taxane binding is a two-step process ^33^. The first step involves initial low affinity binding to the outer wall of MTs near pore type I sites, which then allows translocation of the drug to its high affinity, but kinetically unfavorable, luminal binding site (Supplementary Figure 6A).

Using different chemical probes that interact with the outer *versus* inner MT surface, we treated native cytoskeletons from sensitive and resistant cells expressing canonical only *versus* both CLIP-170 variants. Treatment with hexaflutax, a fluorescently labelled taxoid known to bind to the outer-only site of the MT pore^34^, revealed significantly reduced drug binding to native cytoskeletons from resistant *versus* sensitive cell lines (p < 0.0001) (Figure 4A).

On the other hand, cyclostreptin is known to bind covalently to both the MT pores and luminal taxoid binding site ^33,35^ and strongly inhibits taxane binding. We quantified Flutax-2 binding in cyclostreptin pre-treated native cytoskeleton from sensitive and resistant GC cell lines (Figure 4B). We observed significant reduction in Flutax-2 binding proportional to the duration of cyclostreptin pre-treatment in sensitive TMK1 (0 vs 2 min; p < 0.0001) but not in the resistant Hs746T cells (Figure 4C). In addition, CLIP-170-KD had no effect in TMK1 cells following KD of full-length only, while it significantly reduced Flutax-2 binding in cyclostreptin pre-treated Hs746T cells following KD of both canonical and short variant (Supplementary Figure 7). Together these results suggested that CLIP-170S in Hs746T cells likely obstructs the MT pore, thus kinetically impairing cyclostreptin’s irreversible binding to the site and therefore preventing it from abolishing Flutax-2 binding.

To further probe the involvement of the MT pore, we used peloruside, a compound that binds to a distinct non-taxoid pocket on tubulin ^36,37^ and does not interact with the MT pore ^38^. We found that peloruside was equally effective in both taxane sensitive and resistant GC cells, including the isogenic Hs746T cells stably expressing CLIP-170 or scrambled shRNA (Supplementary Figure 8). Taken together these data strongly support a mechanism whereby CLIP-170S obstructs the MT pore and impairs effective drug target engagement (working model, Supplementary Figure 6B).

### Computational analysis predicts Imatinib as a top candidate to reverse taxane resistance in GC by selective depletion of CLIP-170S

We have previously described how different data types can be combined in a single computational framework to identify protein targets for small molecules in development and to predict new molecules to overcome drug resistance ^39^. Here we used gene expression signatures, derived following RNAseq expression analysis of untreated *vs* DTX-treated sensitive and resistant cell lines, and generated a DTX-resistance signature for our panel of GC cell lines. We then evaluated a set of FDA-approved drugs and active compounds and ranked them based on their potential to reverse this DTX-resistance signature. This analysis identified imatinib as the top hit compound predicted to reverse this taxane resistance in GC cells (Figure 5A). Imatinib is a BCR-Abl inhibitor FDA approved for the treatment of chronic myelogenous leukemia ^40,41^ without any mechanistic implications in MT biology or taxane activity. While treatment with imatinib alone did not show differential activity in sensitive *versus* resistant cells, when combined with DTX it completely reversed taxane resistance in Hs746T and other resistant cell lines (Figure 5B and data not shown) strongly indicating a mechanistic interaction between the two drugs. Intriguingly, we demonstrate that a mere 3 hr treatment with imatinib lead to selective and profound inhibition of CLIP-170S in Hs746T without affecting canonical CLIP-170 in neither resistant nor sensitive GC cells (Figure 5C). Importantly, the imatinib concentration that depletes CLIP-170S and restores taxane sensitivity, is a physiologically achievable concentration in patients ^42^.

To further determine if Imatinib-mediated reversal of taxane resistance was specific to CLIP-170S inhibition we performed a cytotoxicity experiment in Hs746T-CLIP-170-KD. Imatinib did not further sensitize these cells to DTX, confirming the central role of CLIP-170S in this synergistic interaction (Figure 5D). These data were corroborated by combination index analysis to assess the extent of synergy between the two drugs, which showed synergistic interaction in the Hs746T cells (CI ∼0.5) but lack thereof upon CLIP-170 depletion (CI> 1.2) (Supplementary Figure 9).

### CLIP-170S expression is highly prevalent in tumor biopsies from GC patients

To investigate the clinical relevance of CLIP-170S expression, we assessed CLIP-170S expression in a cohort of GC patients. We identified the presence of CLIP-170S in 25/37 tumor samples, giving a prevalence of CLIP-170S of 67.5% (95% CI 52.2 – 82.8 %) (data not shown).

## Discussion

The microtubule cytoskeleton is arguably one of the most validated therapeutic target in oncology, as evidenced by the use of taxanes in a broad spectrum of tumors. Taxanes induce cell death by binding to and stabilizing MTs, thus, blocking essential MT-dependent processes including signaling, trafficking and mitosis ^43^ and reviewed in ^44,45^. Although widely used, the clinical success of taxane therapy has been marred by intrinsic and acquired resistance, the molecular underpinnings of which have remained elusive.

Here we identify a previously unrecognized variant of the plus-end binding protein CLIP-170, which we named CLIP-170S, as it lacks the first 150 amino acids and results in a shorter protein of about 140 kD. The missing N-terminal serine-rich domain and the first Cap-Gly domain, are required for effective MT plus-end tracking ^31,32^, thus, resulting in mislocalization of CLIP-170S from the MT ends to the MT lattice (Figure 3).

Using a panel of GC cell lines and patient tumors we show, for the first time, that CLIP-170S mislocalization to the MT lattice results in taxane resistance by occlusion of the MT pore which limits taxane access to its high affinity binding site in the MT lumen (Figure 3 and 4). To the best of our knowledge the MT pore has not been previously implicated in taxane resistance *in vitro*, which together with the high prevalence of CLIP-170S in gastric tumors, provides a paradigm shift in our understanding of clinical taxane resistance.

The discovery of the MT pore type I as an external, transient interaction site for taxanes on the outer MT wall helped explain how taxanes reach the kinetically unfavorable luminal binding site ^33,34,46^ and reviewed in ^47^. Nevertheless, there has been paucity of data regarding biological roles for the MT pore beyond facilitating taxane internalization to the MT lumen ^22^. Our work provides new insights where obstruction of the MT pore alone diminishes taxane activity, consistent with absence of tubulin structural alterations in our model, thus representing a never-before described mechanism of resistance.

Having identified the integrity of the MT pore as an actionable therapeutic target, we employed systems biology and discovered that Imatinib (Gleevec) reverses taxane resistance in CLIP-170S expressing GC by restoring the MT pore integrity following selective depletion of CLIP-170S. This was an unexpected finding as Imatinib primarily targets BCR-ABL fusion protein in chronic mylogenous leukemia (CML)^40,41,48,49^ as well as other receptor tyrosine kinases (RTKs) including PDGFR, VEGFR, c-Abl and c-kit^50,51^, none of which was expressed in our GC cell line panel (data not shown). We envision that this work will pave the way for unprecedented clinical benefits for taxane-refractory patients expressing CLIP-170S or potentially other MT-pore occluding proteins.

---

<u>**Supplementary Figure 1:**</u> A) Representative pictures of Flutax-2-labeled native cytoskeletons from a DTX-resistant cell line SCH. bar = 20 µm. B) Box-plot representation of Flutax-2 fluorescence intensity in SCH. 5-95% confidence interval graphs are shown (n = 20 – 70 cells/time point/cell line), statistical values between 0 and 60 min are shown; Mann-Whitney test.

<u>**Supplementary Figure 2:**</u> Graphical representation of taxane binding kinetics to native-cytoskeletons from sensitive (TMK1) and resistant (Hs746T) cell lines. A) For *k*_*on*_ measurements, cells were treated with 1µM Flutax-2 for 10s, 15s, 30s, 60s and 80s. Following washout, cells were imaged using a spinning disk confocal microscope. Increasing fluorescence intensities corresponding to increasing Flutax-2 incubation times were then used to calculate *k*_*on*_ values in both TMK1 and Hs746T cells. B) For *k*_*off*_ measurements, cells were treated with Flutax-2, followed by replacement of Flutax-2 with unlabeled DTX (0s). Flutax-2 fluorescence intensity was recorded at 0, 30, 60, 120, 180 and 240s. Decreasing fluorescence intensities at different time points were then used to calculate *k*_*off*_ values in both TMK1 and Hs746T cells. Results are representative of two independent biological repeats.

<u>**Supplementary Figure 3:**</u> 5’RACE reveals presence of an alternative CLIP-170 transcript starting in exon 3, in Hs746T cells. Experiment performed using RNA extracted from DTX-resistant Hs746T cells. 5’RACE fragments were cloned in a pRACE plasmid and individual clones were sequenced to analyze CLIP-170 transcripts. A) Representative gel showing nine different clones, eight of which contain the canonical CLIP-170 transcript starting in exon 1. One clone (clone 8) contains an alternate transcript starting in the middle of exon 3. B) Representative sequencing results from clones 2 and 8 are shown. Canonical transcription (Exon1) and translation (Exon2) start sites for clone 2 and putative translation start site (Exon 3) for clone 8 are indicated using arrows. C) Schematic representation comparing the exon structure of CLIP-170 versus CLIP-170S as elucidated from 5’RACE experiment in (A) and (B). Arrows indicate position of canonical translation start site for CLIP-170 and putative translation start site for CLIP-170S.

<u>**Supplementary Figure 4:**</u> Immunofluorescence staining using tubulin and GFP antibodies was performed in COS-7 cells ectopically expressing either the GFP-tagged CLIP-170 (CLIP-170) or GFP-tagged CLIP-170S (CLIP-170S). Similar to live-cell imaging results (Figure 3B), immunofluorescence staining of fixed cells using the GFP antibody also revealed a comet-like pattern of canonical CLIP-170 in contrast to the MT lattice distribution pattern exhibited by CLIP-170S. Notice the near complete overlap between GFP and tubulin staining in cells expressing CLIP-170S only.

<u>**Supplementary Figure 5:**</u> Flutax-2 residence time on MTs is restored upon depletion of CLIP-170S in Hs746T-CLIP-170-KD cells. Box-plot representation of Flutax-2 fluorescence intensity in Hs746T-scrambled (*top*) and Hs746T-CLIP-170-depleted resistant cells (*bottom*). 5-95% confidence intervals graphs are shown; statistical values for each cell line between 0 and 45 min are shown; Mann-Whitney test; n.s.; not significant. DTX sensitivity data for Hs746T-scrambled and Hs746T-CLIP-170-KD are shown in Figure 3D.

<u>**Supplementary Figure 6:**</u> Working model of CLIP-170S mediated taxane resistance A) Canonical CLIP-170 binds at MT plus-ends (cell cartoon to the right) and does not affect taxane binding, depicted here as a 2-step process. Step 1: taxane first binds to the low affinity binding site on the MT pore surface (Step 1), gets internalized through the pore and then binds to its high affinity binding site in the MT lumen (Step 2). B) CLIP-170S is mislocalized to the MT lattice (cell to the right), obstructing taxane binding to the MT pore (Step 1) which then limits access of taxane into the MT lumen (Step 2), thus, rendering cells taxane-resistant.

<u>**Supplementary Figure 7:**</u> Fluorescence intensity of Flutax-2 binding to native cytoskeletons pre-incubated with cyclostreptin in TMK1-CLIP-170-KD and Hs746T-CLIP-170-KD cells. Significant loss in Flutax-2 binding was observed in both cell lines following a 2 min cyclostreptin pre-incubation Mann-Whitney test; 5-95% confidence interval shown. Notice the significant drop in Flutax-2 binding in Hs746T-CLIP-170-KD cells, in contrast to the results shown in Figure 4C where cyclostreptin fails to abolish Flutax-2 binding in Hs746T cells expressing both canonical and short CLIP variants.

<u>**Supplementary Figure 8:**</u> Cytotoxicity assays of sensitive (TMK1 and AZ521) and resistant (Hs746T) cells and its derivative Hs746T-scrambled and Hs746T-CLIP-170-KD cell lines are shown. Treatment with DTX (A) or Peloruside (B) for 72 hr. Peloruside, which does not traverse through the MT pore, is equally effective in all cell lines in contrast to DTX. Table with representative IC_50_ values is shown. Results are representative of three biological repeats.

<u>**Supplementary Figure 9:**</u> Combination index analysis reveals synergistic drug interaction between DTX and IMA in Hs746T-scramble cells (A) in contrast to the Hs746T-CLIP-170-KD cells (B). In this analysis Combination Index value of less than 1 (CI < 1) indicates synergisms while CI>1 indicates antagonism. Notice the profound drug synergy in Hs746T cells (CI∼ 0.5) but not in CLIP-170 KD cell line (CI > 1.2), suggesting synergistic effects of the two drugs only in cell line specifically expressing CLIP-170S protein.

<u>**Supplementary movies 1 and 2:**</u> Live cell imaging of COS-7 cells with ectopic expression of similarly low levels of either GFP-tagged CLIP-170 (movie 1) or CLIP-170S (movie 2). 1) Similar to the snapshot indicated in Figure 3B, GFP-tagged canonical CLIP-170 shows a comet like distribution indicative of plus-end localization. 2) Two different cells overexpressing high (left) vs low (right) levels of GFP-tagged CLIP-170S can be seen. CLIP-170S decorates the MT lattice even in cells expressing low levels of the protein (right), similar to the pattern expected in cells expressing high levels of GFP-tagged CLIP-170S (left).

## Acknowledgements

This work was supported in part by the NIH T32 Training grant 5T32CA062948 (to KK), the NIH T32 Training grant 5T32CA062948 (to GG), by Clinical and Translational Science Center at Weill Cornell Medicine NIH/NCATS grant ULTR00457 (to GG), the NIH/NCI R01CA228512 (to P.G., M.A.S. and O.E.), NIH/NCI R21 CA216800 (to P.G), NIH/NCI R01 CA179100 (to P.G) and by the and Ministerio de Economía y Competitividad grant BFU2016-75319-R and European Union H2020-MSCA-ITN-ETN/0582 ITN TUBINTRAIN (Awarded to J.F.D.). D.S. was supported by the Intramural Research Program of the Eunice Kennedy Shriver National Institute of Child Health and Human Development, NIH.

## Author contributions

K.K., P.V.T., M.A.S., and P.G. conceptualized and designed the study, analyzed data and wrote the manuscript. K.K., P.V.T., G.G., E.V.N. performed experiments and analyzed data. N.M., O.E. performed computational analysis and reviewed and edited the manuscript. H.V.G, D.S., I.B., J.F.D., N.M., O.E. designed experiments, provided comments, analyzed data, reviewed and edited the manuscript.

## Author information

Drs. Neel Madhukar and Olivier Elemento are the co-founders and equity stakeholders of OneThree Biotech, an artificial intelligence-driven drug discovery and development company. All other authors have no competing interests and have nothing to declare.

## Data availability

The data that support this study are available from the corresponding authors upon request.

