## Supplementary Figures for "Imatinib overrides taxane resistance by selective inhibition of novel CLIP1 variant obstructing the microtubule pore"

Supplementary Figure 1

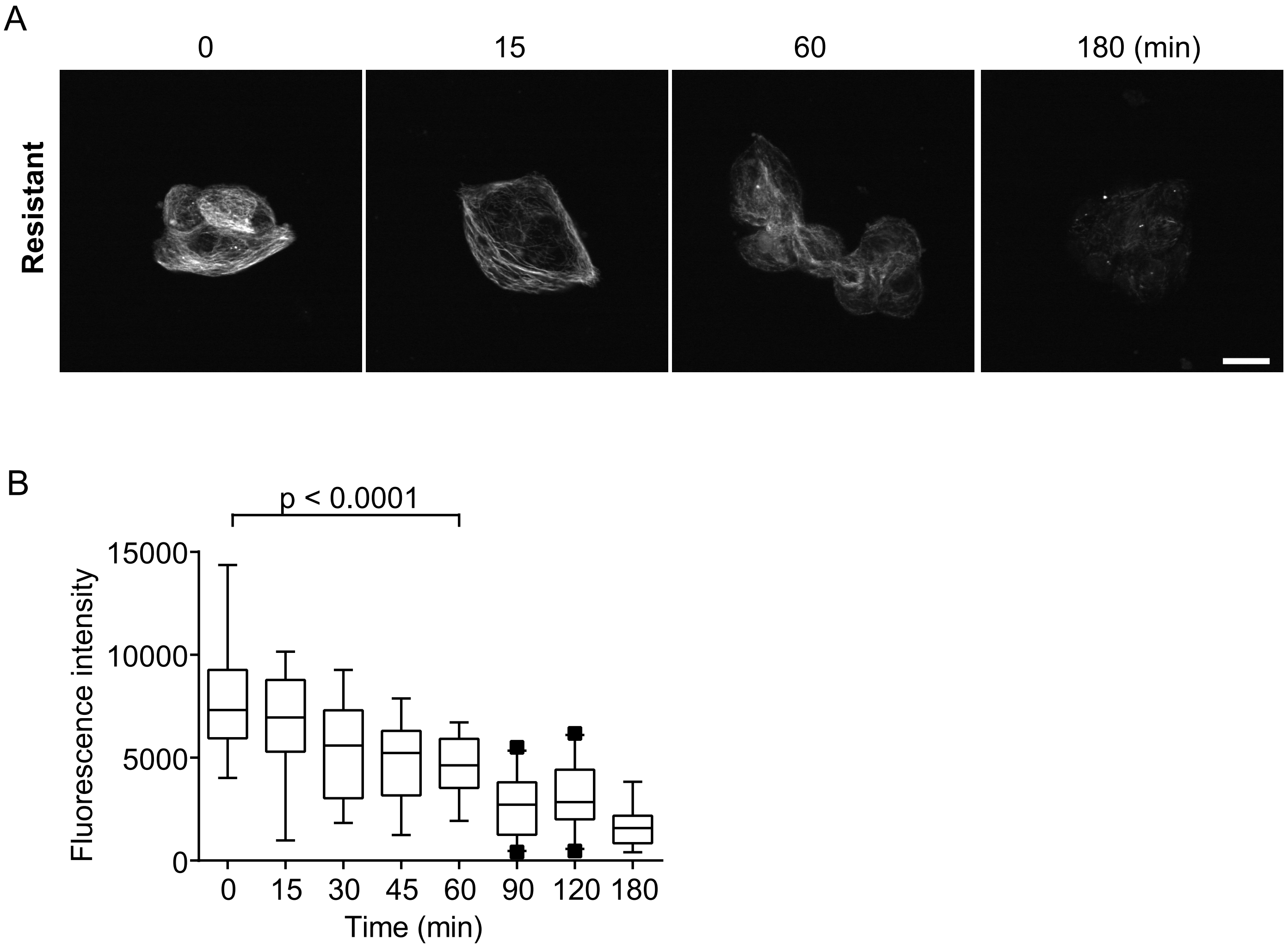


**Supplementary Figure 1:** A) Representative pictures of Flutax-2-labeled native cytoskeletons from a DTX-resistant cell line SCH. bar = 20 µm. B) Box-plot representation of Flutax-2 fluorescence intensity in SCH. 5-95% confidence interval graphs are shown (n = 20 - 70 cells/time point/cell line), statistical values between 0 and 60 min are shown; Mann-Whitney test.

### Supplemental Figure 2

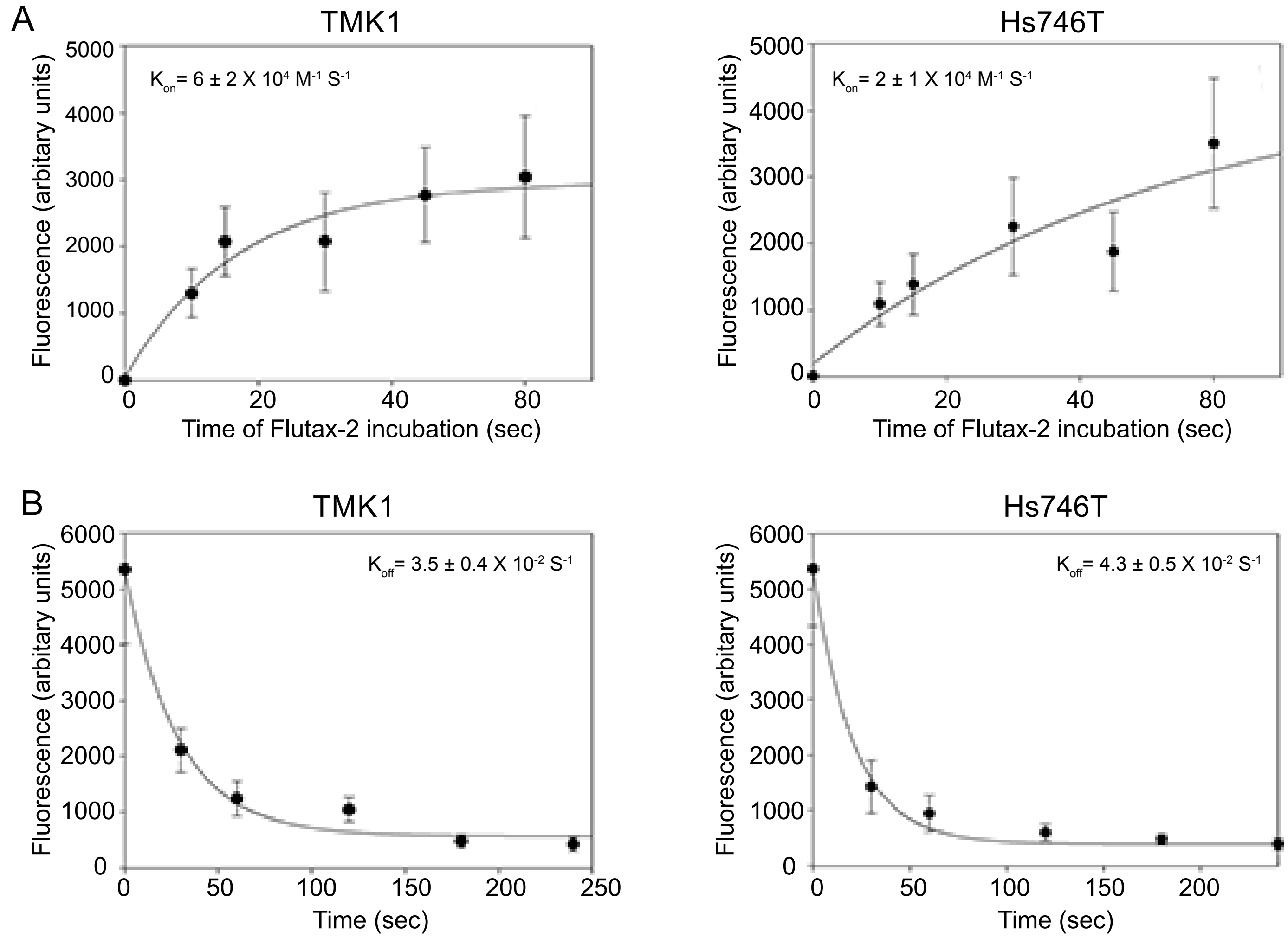


**Supplementary Figure 2:** Graphical representation of taxane binding kinetics to native-cytoskeletons from sensitive (TMK1) and resistant (Hs746T) cell lines. A) For $k_{on}$ measurements, cells were treated with 1μM Flutax-2 for 10s, 15s, 30s, 60s and 80s. Following washout, cells were imaged using a spinning disk confocal microscope. Increasing fluorescence intensities corresponding to increasing Flutax-2 incubation times were then used to calculate $k_{on}$ values in both TMK1 and Hs746T cells. B) For $k_{off}$ measurements, cells were treated with Flutax-2, followed by replacement of Flutax-2 with unlabeled DTX (0s). Flutax-2 fluorescence intensity was recorded at 0, 30, 60, 120, 180 and 240s. Decreasing fluorescence intensities at different time points were then used to calculate $k_{off}$ values in both TMK1 and Hs746T cells. Results are representative of two independent biological repeats.

### Supplementary Figure 3

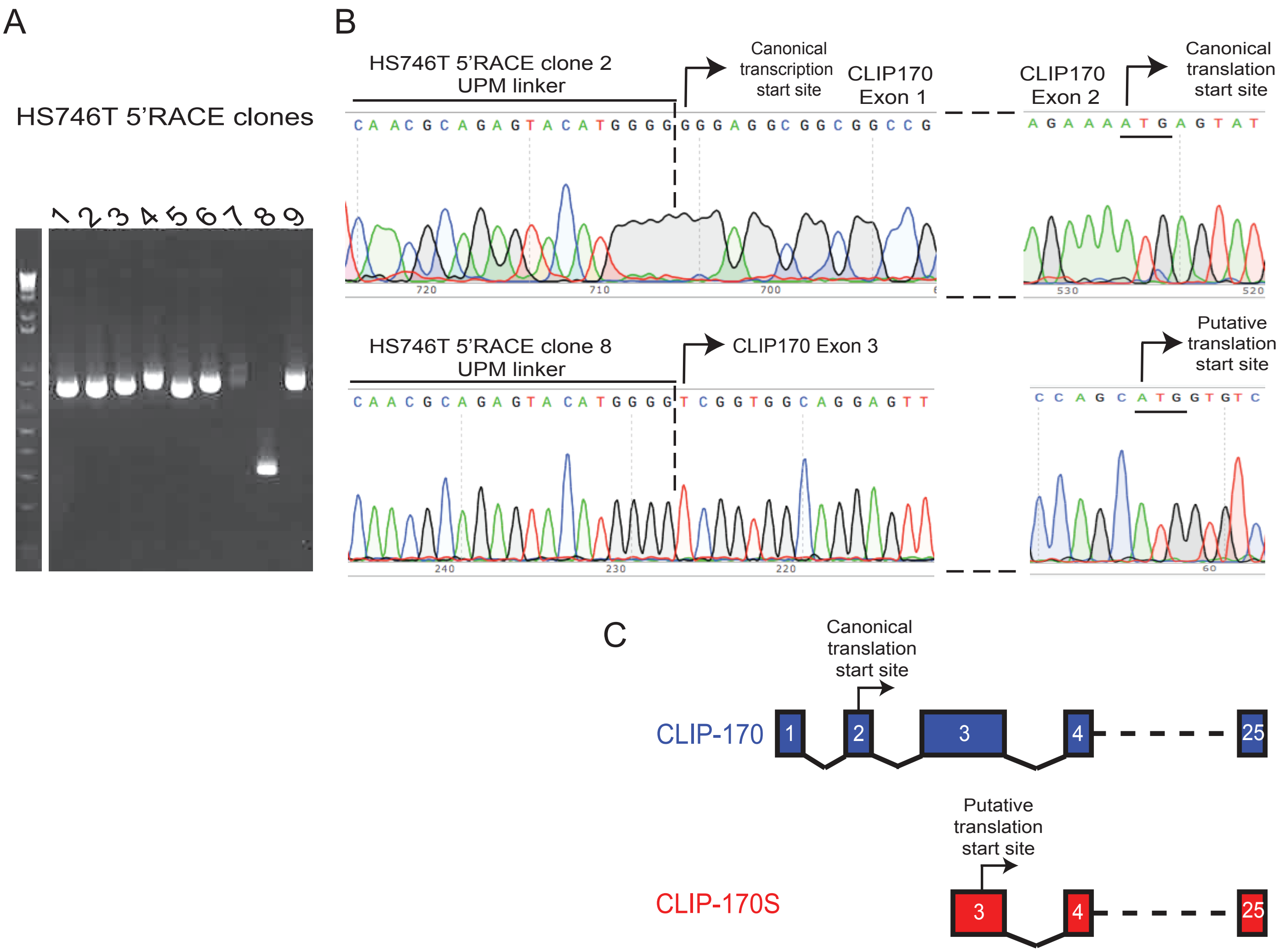


**Supplementary Figure 3:** 5'RACE reveals presence of an alternative CLIP-170 transcript starting in exon 3, in Hs746T cells. Experiment performed using RNA extracted from DTX-resistant Hs746T cells. 5'RACE fragments were cloned in a pRACE plasmid and individual clones were sequenced to analyze CLIP-170 transcripts. A) Representative gel showing nine different clones, eight of which contain the canonical CLIP-170 transcript starting in exon 1. One clone (clone 8) contains an alternate transcript starting in the middle of exon 3. B) Representative sequencing results from clones 2 and 8 are shown. Canonical transcription (Exon1) and translation (Exon2) start sites for clone 2 and putative translation start site (Exon 3) for clone 8 are indicated using arrows. C) Schematic representation comparing the exon structure of CLIP-170 versus CLIP-170S as elucidated from 5'RACE experiment in (A) and (B). Arrows indicate position of canonical translation start site for CLIP-170 and putative translation start site for CLIP-170S.

Supplementary Figure 4

A)

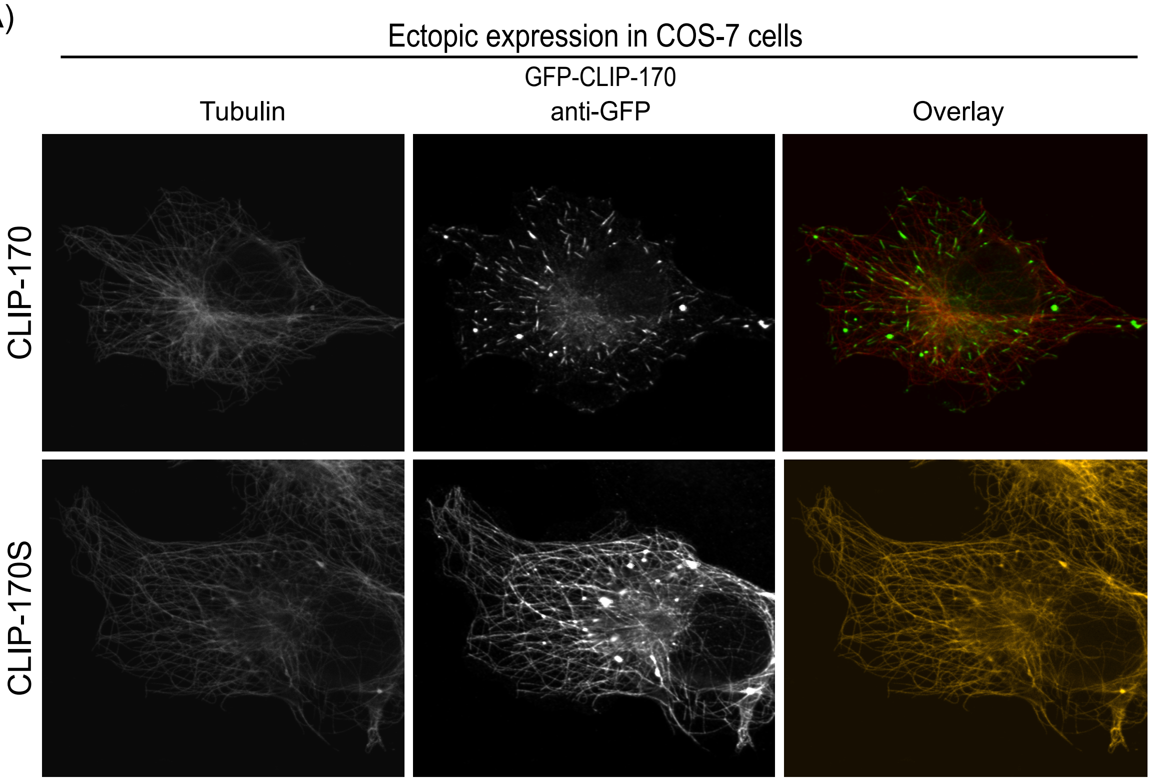


**Supplementary Figure 4:** Immunofluorescence staining using tubulin and GFP antibodies was performed in COS-7 cells ectopically expressing either the GFP-tagged CLIP-170 (CLIP-170) or GFP-tagged CLIP-170S (CLIP-170S). Similar to live-cell imaging results (Figure 3B), immunofluorescence staining of fixed cells using the GFP antibody also revealed a comet-like pattern of canonical CLIP-170 in contrast to the MT lattice distribution pattern exhibited by CLIP-170S. Notice the near complete overlap between GFP and tubulin staining in cells expressing CLIP-170S only.

Supplementary Figure 5

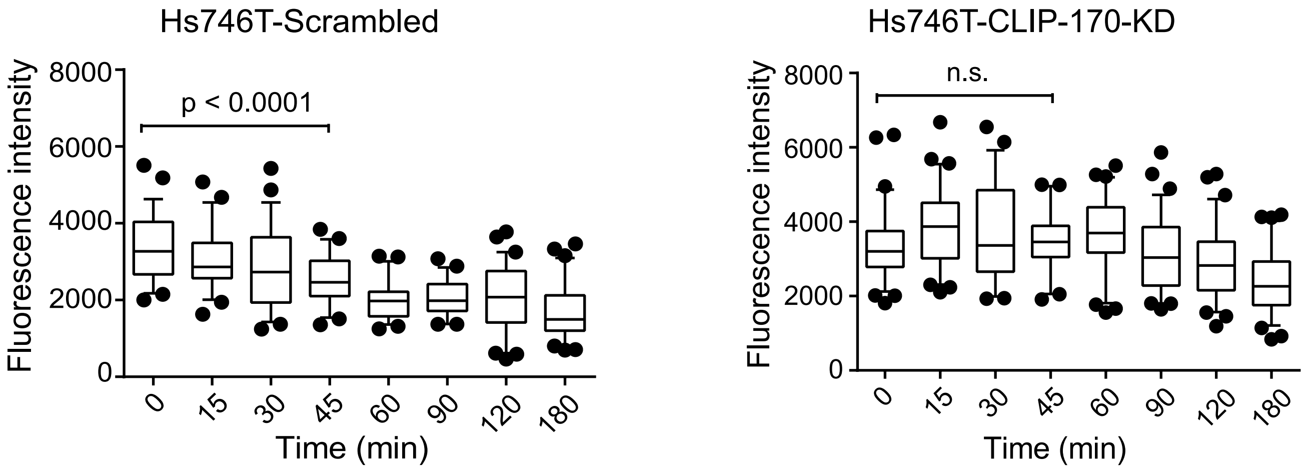


**Supplementary Figure 5:** Flutax-2 residence time on MTs is restored upon depletion of CLIP-170S in Hs746T-CLIP-170-KD cells. Box-plot representation of Flutax-2 fluorescence intensity in Hs746T-scrambled (top) and Hs746T-CLIP-170-depleted resistant cells (bottom). 5-95% confidence intervals graphs are shown; statistical values for each cell line between 0 and 45 min are shown; Mann-Whitney test; n.s.; not significant. DTX sensitivity data for Hs746T-scrambled and Hs746T-CLIP-170-KD are shown in Figure 3D.

### Supplementary Figure 6

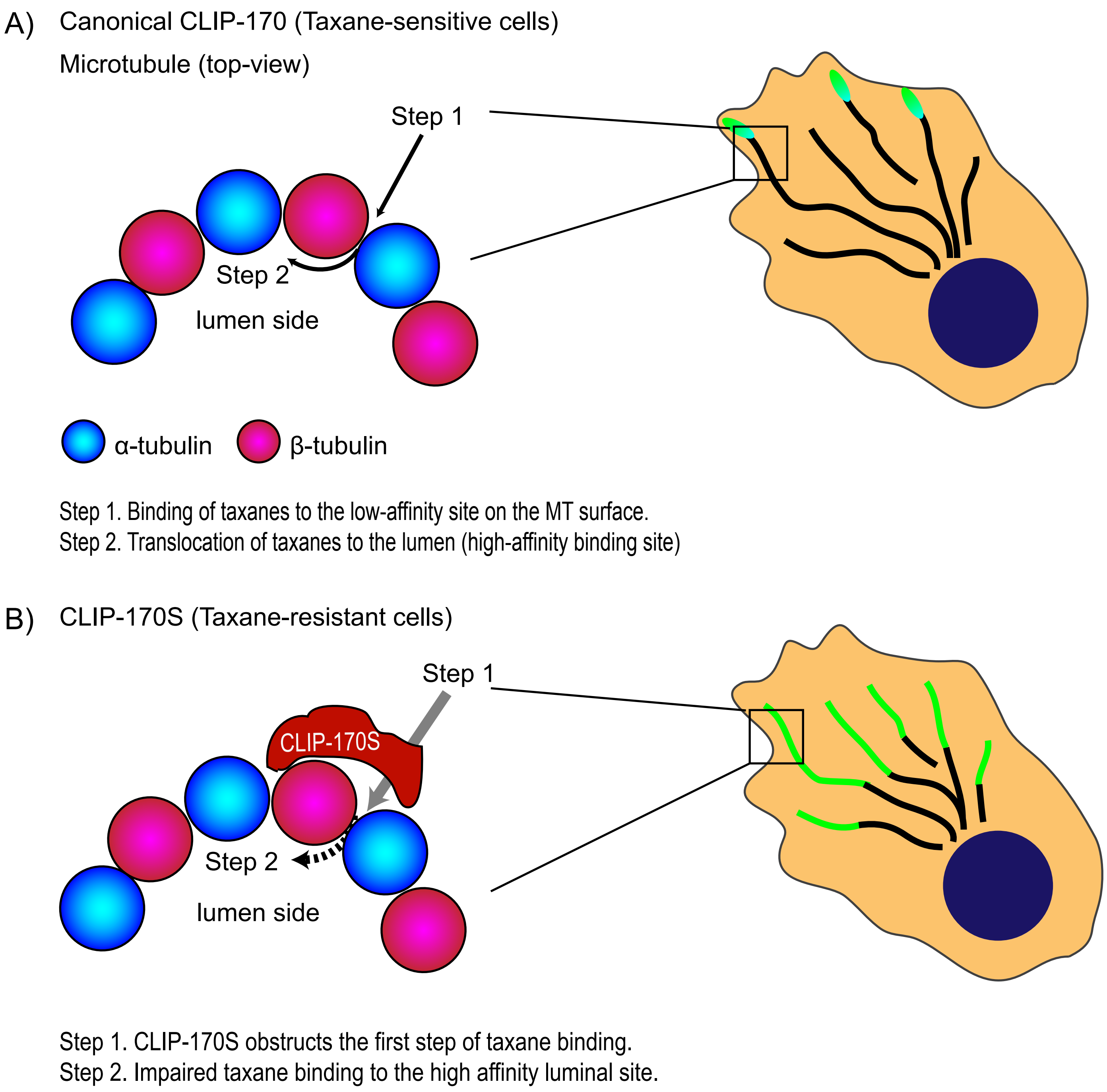


**Supplementary Figure 6:** Working model of CLIP-170S mediated taxane resistance A) Canonical CLIP-170 binds at MT plus-ends (cell cartoon to the right) and does not affect taxane binding, depicted here as a 2-step process. Step 1: taxane first binds to the low affinity binding site on the MT pore surface (Step 1), gets internalized through the pore and then binds to its high affinity binding site in the MT lumen (Step 2). B) CLIP-170S is mislocalized to the MT lattice (cell to the right), obstructing taxane binding to the MT pore (Step 1) which then limits access of taxane into the MT lumen (Step 2), thus, rendering cells taxane-resistant.

### Supplementary Figure 7

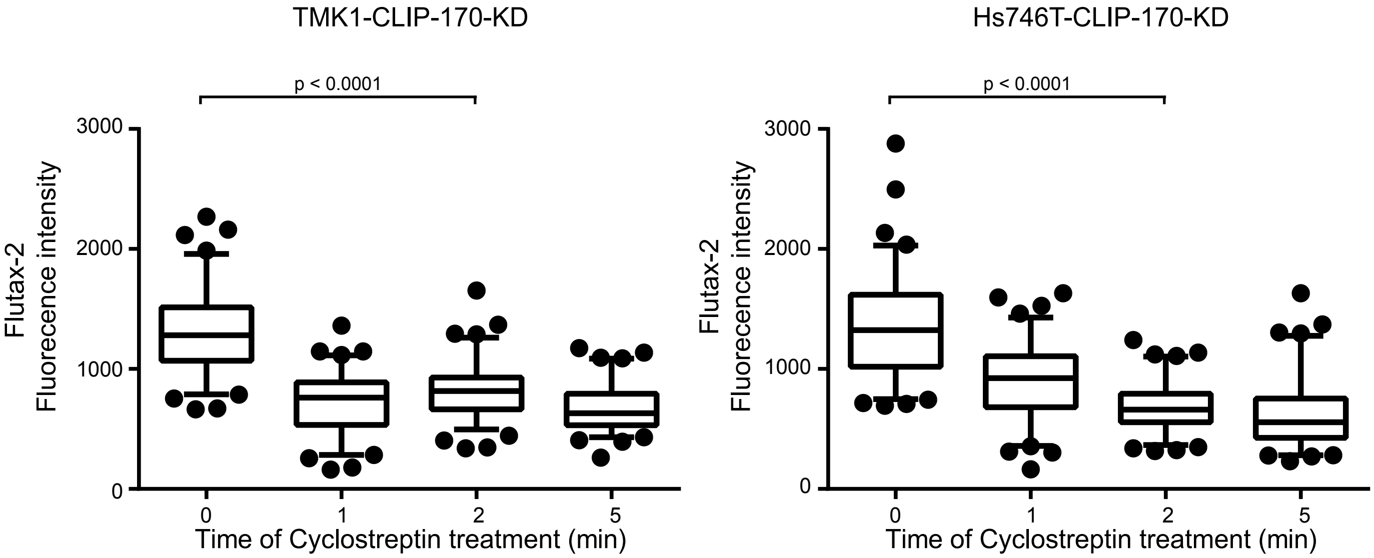


**Supplementary Figure 7:** Fluorescence intensity of Flutax-2 binding to native cytoskeletons pre-incubated with cyclostreptin in TMK1-CLIP-170-KD and Hs746T-CLIP-170-KD cells. Significant loss in Flutax-2 binding was observed in both cell lines following a 2 min cyclostreptin pre-incubation Mann-Whitney test; 5-95% confidence interval shown. Notice the significant drop in Flutax-2 binding in Hs746T-CLIP-170-KD cells, in contrast to the results shown in Figure 4C where cyclostreptin fails to abolish Flutax-2 binding in Hs746T cells expressing both canonical and short CLIP variants.

Supplementary figure 8

A B

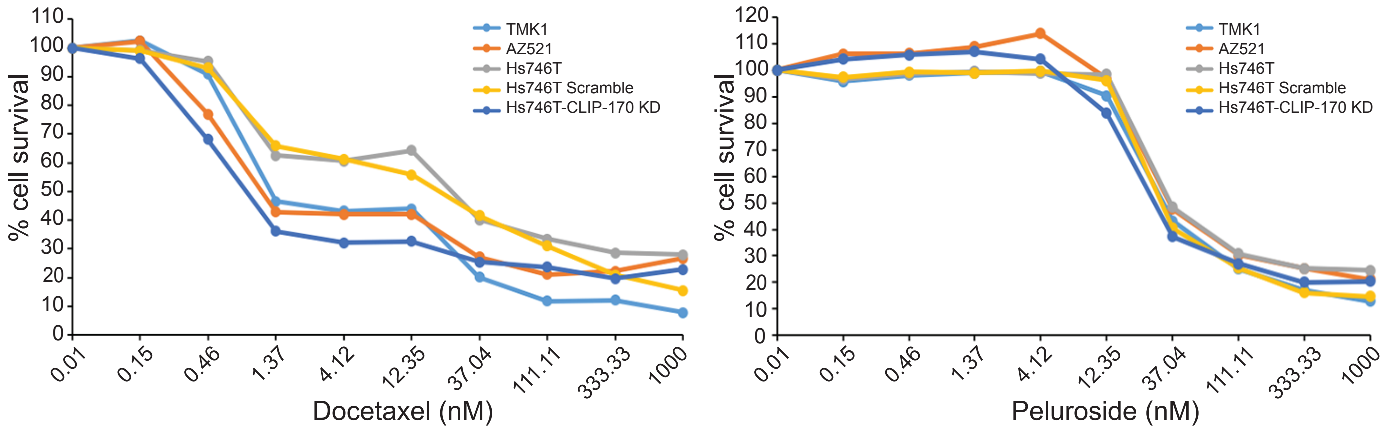


| | $IC_{50}$ DTX | $IC_{50}$ Peluroside |
|---|---|---|
| TMK1 | 1.30 nM | 33.50 nM |
| AZ521 | 1.18 nM | 35.85 nM |
| Hs746T | 26.92 nM | 36.25 nM |
| Hs746T Scrambled | 22.37 nM | 32.82 nM |
| Hs746T-CLIP-170 KD | 0.97 nM | 30.25 nM |

**Supplementary Figure 8:** Cytotoxicity assays of sensitive (TMK1 and AZ521) and resistant (Hs746T) cells and its derivative Hs746T-scrambled and Hs746T-CLIP-170-KD cell lines are shown. Treatment with DTX (A) or Peloruside (B) for 72 hr. Peloruside, which does not traverse through the MT pore, is equally effective in all cell lines in contrast to DTX. Table with representative $IC_{50}$ values is shown. Results are representative of three biological repeats.

Supplementary Figure 9

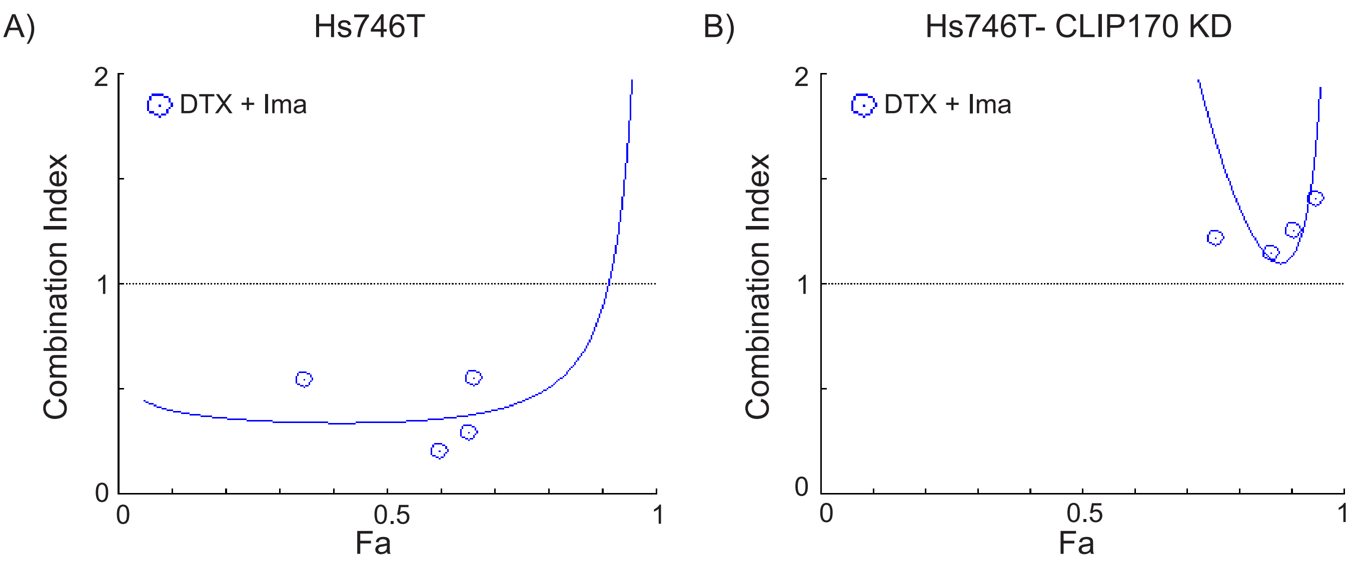


**Supplementary Figure 9:** Combination index analysis reveals synergistic drug interaction between DTX and IMA in Hs746T-scramble cells (A) in contrast to the Hs746T-CLIP-170-KD cells (B). In this analysis Combination Index value of less than 1 (CI < 1) indicates synergisms while CI>1 indicates antagonism. Notice the profound drug synergy in Hs746T cells (CI~ 0.5) but not in CLIP-170 KD cell line (CI > 1.2), suggesting synergistic effects of the two drugs only in cell line specifically expressing CLIP-170S protein.
